# Comparative Analysis of Single-Cell RNA Sequencing Methods

**DOI:** 10.1101/035758

**Authors:** Christoph Ziegenhain, Beate Vieth, Swati Parekh, Björn Reinius, Martha Smets, Heinrich Leonhardt, Ines Hellmann, Wolfgang Enard

## Abstract

**Background:** Single-cell RNA sequencing (scRNA-seq) offers exciting possibilities to address biological and medical questions, but a systematic comparison of recently developed protocols is still lacking.

**Results:** We generated data from 447 mouse embryonic stem cells using Drop-seq, SCRB-seq, Smart-seq (on Fluidigm C1) and Smart-seq2 and analyzed existing data from 35 mouse embryonic stem cells prepared with CEL-seq. We find that Smart-seq2 is the most sensitive method as it detects the most genes per cell and across cells. However, it shows more amplification noise than CEL-seq, Drop-seq and SCRB-seq as it cannot use unique molecular identifiers (UMIs). We use simulations to model how the observed combinations of sensitivity and amplification noise affects detection of differentially expressed genes and find that SCRB-seq reaches 80% power with the fewest number of cells. When considering cost-efficiency at different sequencing depths at 80% power, we find that Drop-seq is preferable when quantifying transcriptomes of a large numbers of cells with low sequencing depth, SCRB-seq is preferable when quantifying transcriptomes of fewer cells and Smart-seq2 is preferable when annotating and/or quantifying transcriptomes of fewer cells as long one can use in-house produced transposase.

**Conclusions:** Our analyses allows an informed choice among five prominent scRNA-seq protocols and provides a solid framework for benchmarking future improvements in scRNA-seq methodologies.

## Background

Genome-wide quantification of mRNA transcripts can be highly informative for the characterization of cellular states and to understand regulatory circuits and processes^1,2^. Ideally, such data are collected with high spatial resolution, and scRNA-seq now allows for transcriptome-wide analyses of individual cells, revealing new and exciting biological and medical insights^3–5^. scRNA-seq requires the isolation of single cells and the conversion of their RNA into cDNA libraries that can be quantified using high-throughput sequencing^4,6^. How well single-cell transcriptomes can be characterized depends on many factors, including the sensitivity of the method, i.e. which and how many mRNAs can be detected, its accuracy, i.e. how well the quantification corresponds to the actual concentration of mRNAs and its precision, i.e. with how much technical noise mRNAs are quantified. Of high practical relevance is also the efficiency of the method, i.e. the monetary cost to characterize single cells, e.g. at a certain level of precision. In order to make a well-informed choice among available scRNA-seq methods, it is important to estimate these parameters comparably. Each method is likely to have its own strengths and weaknesses. For example, it has previously been shown that scRNA-seq conducted in the small volumes available in the automated microfluidic platform from Fluidigm (Smart-seq protocol on the C1-platform) performs better than Smart-seq or other commercially available kits in microliter volumes^7^. Furthermore, the Smart-seq protocol has been optimized for sensitivity, even full-length coverage, accuracy and cost^8^ and this improved “Smart-seq2” protocol^9^ has also become widely used^10–14^.

Other protocols have sacrificed full-length coverage for 3’ or 5’ sequencing of mRNAs in order to sequence part of the primer used for cDNA generation. This enables early barcoding of libraries, i.e. the incorporation of well-specific or cell-specific barcodes, allowing to multiplex cDNA amplification and library generation and thereby increasing the throughput of scRNA-seq library generation by one to three orders of magnitude^15–19^.

Additionally, this approach allows the incorporation of Unique Molecular Identifiers (UMIs), random nucleotide sequences that tag individual mRNA molecules and hence allow for the distinction between original molecules and amplification duplicates that derive from the cDNA or library amplification^18,20,21^. Utilization of UMI information leads to improved quantification of mRNA molecules^22,23^ and has been implemented in several scRNA-seq protocols, such as STRT^22^, CEL-seq^23^, Drop-seq^17^, inDrop^19^, MARS-seq^16^ or SCRB-seq^15^. However, a thorough and systematic comparison of scRNA-seq methods, evaluating sensitivity, accuracy, precision and efficiency is still lacking. To address this issue, we analyzed 482 scRNA-seq libraries from mouse embryonic stem cells (mESCs), generated using five different methods with two technical replicates for each method (Figure 1).

**Fig. 1.**
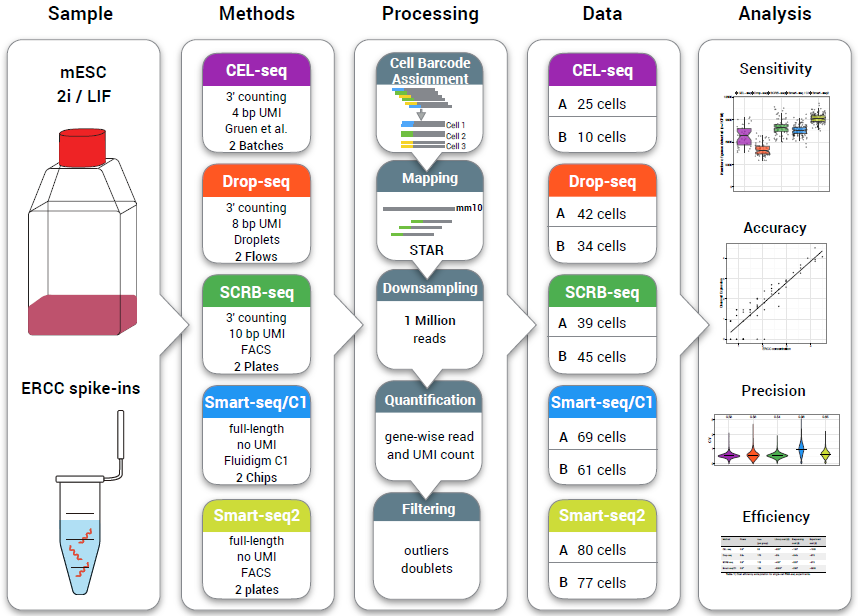
| Schematic overview of the experimental and computational workflow. Mouse embryonic stem cells (mESCs) cultured in 2i/LIF and ERCC spike-in RNA were used to generate single-cell RNA-seq data with five different library preparation methods (CEL-seq, Drop-seq, SCRB-seq, Smart-seq/C1 and Smart-seq2). The methods differ in the usage of unique molecular identifier sequences (UMI), which allow the discrimination between reads derived from original mRNA molecules and duplicates during cDNA amplification. Data processing was identical across methods and analyzed cell numbers per method and replicate are given, which were used to compare sensitivity, accuracy, precision and cost-efficiency. The five scRNA-seq methods are denoted by color throughout the figures of this study: purple – CEL-seq, orange – Drop-seq, green SCRB-seq, blue – Smart-seq, yellow – Smart-seq2.

## Results

### Generation of scRNA-seq libraries

We generated scRNA-seq libraries from mouse embryonic stem cells (mESCs) in two independent replicates using Smart-seq^24^, Smart-seq2^8^, Drop-seq^17^ and SCRB-seq^15^. Additionally, we used available scRNA-seq data^23^ from mESCs that was generated using CEL-seq^18^. An overview of the employed methods and their library generation workflows is provided in (Figure 2) and in Supplementary Table 1.

**Fig. 2.**
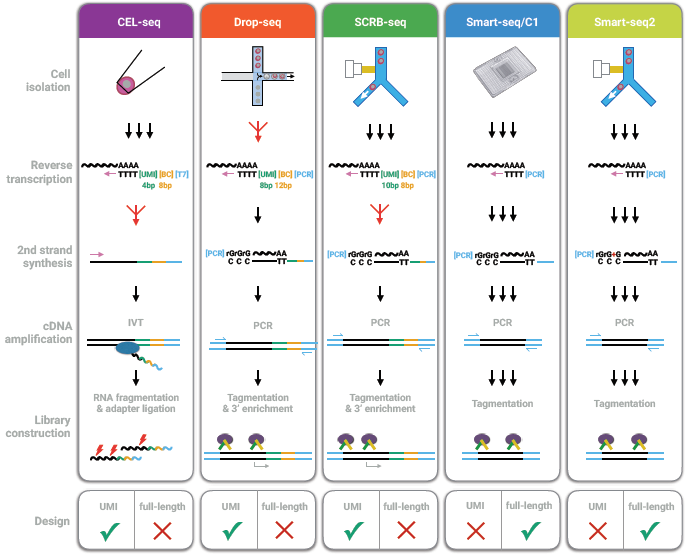
| Schematic overview of library preparation steps. For details see text.

For each replicate of the Smart-seq protocol, we performed a run on the C1 platform from Fluidigm (Smart-seq/C1) using the 10-17 μm mRNA-seq Integrated Fluidic Circuit (IFCs) microfluidic chips that can automatically capture up to 96 cells^7^. We imaged the cells to identify doublets (see below) and added lysis buffer together with External RNA Control Consortium spike-ins (ERCCs) that consist of 92 poly-adenylated synthetic RNA transcripts spanning a range of concentrations^25^. We used the commercially available Smart-seq kit (Clontech) that uses oligo-dT priming, template switching and PCR amplification to generate full-length double-stranded cDNA. We harvested the amplified cDNAs and converted them into 96 different sequenceable libraries by tagmentation (Nextera, Illumina) and PCR amplification using indexed primers for multiplexing. Advantages of this system include that single cell isolation and cDNA generation is automated, that captured cells can be imaged, that reaction volumes are small and that full-length cDNA libraries are sequenced.

For each replicate of the Smart-seq2 protocol, we sorted mESCs by flow cytometry into 96-well PCR plates containing lysis buffer and ERCCs. We generated cDNA as described^8,9^ and used an in-house produced Tn5 transposase^26^ to generate 96 libraries by tagmentation. While Smart-seq/C1 and Smart-seq2 are very similar protocols that generate full-length libraries they differ in how cells are isolated, the reaction volume and in that Smart-seq2 has been systematically optimized^8,9^. The main disadvantage of both protocols is that the generation of full-length cDNA libraries precludes an early barcoding step and the incorporation of UMIs.

For each replicate of the SCRB-seq protocol^15^, we also sorted mESCs by flow cytometry into 96-well PCR plates containing lysis buffer and ERCCs. Also similar to Smart-seq2, cDNA is generated by oligo-dT priming, template switching and PCR amplification of full-length cDNA. However, the oligo-dT primers contain well-specific (i.e. cell-specific) barcodes and UMIs. Hence, cDNA from one plate can be pooled and then be converted into RNA-seq libraries, whereas a modified transposon-based fragmentation approach is used that enriches for 3’ ends. The protocol is optimized for small volumes and few handling steps, but it does not generate full-length RNA-seq profiles and its performance compared to other methods is unknown.

The fourth method evaluated was Drop-seq, a recently developed microdroplet-based approach^17^. Similarly to SCRB-seq, each cDNA molecule is labeled with a cell-specific multiplexing barcode and an UMI to count original mRNA molecules. In the case of Drop-seq, over 10^8^ of such barcoded oligo-dT primers are immobilized on beads with each bead carrying a unique cell barcode. A flow of beads suspended in lysis buffer and a flow of a single-cell suspension are brought together in a microfluidic chip that generates nanoliter-sized emulsion droplets. Cells are lysed within these droplets, their mRNA binds to the oligo-dT-carrying beads, and after breaking the droplets, reverse transcription, template switching and library generation is performed for all cells in parallel in a single tube. The ratio of beads to cells (20:1) ensures that the vast majority of the beads have either no (>95% expected when Poisson distributed) or one single cell (4.8% expected) in their droplet and hence ensures that doublets are rare (<0.12% expected)^17^. We benchmarked our Drop-seq setup as recommended^17^ and determined the doublet rate by mixing mouse and human T-cells (~2.5% of sequenced cell transcriptomes; Supplementary (Figure 1a), confirming that the Drop-seq protocol works well in our setup. The main advantage of the protocol is that many scRNA-seq libraries can be generated at low costs. One disadvantage is that the simultaneous inclusion of ERCC spike-ins is not practical for Drop-seq, as their addition would generate cDNA from ERCCs also in all beads that have no cell and hence would approximately double the sequencing costs. As a proxy for the missing ERCC data, we used a published dataset^17^, where ERCC spike-ins were sequenced by the Drop-seq method without single-cell transcriptomes.

Finally, we re-analyzed data^23^ generated using CEL-seq^18^ for which two replicates of scRNA-seq libraries were available for the same cell type and culture conditions (mESCs in 2i/LIF). Similarly to Drop-seq and SCRB-seq, cDNA is tagged with multiplexing barcodes and UMIs. As opposed to the four PCR-based methods described above, CEL-seq relies on linear amplification by in-vitro transcription (IVT) for the initial pre-amplification of singlecell material.

### Processing of scRNA-seq data

For Smart-seq2, Smart-seq/C1, SCRB-seq and Drop-seq we generated libraries from 192, 192, 192 and ~200 cells in the two independent replicates and sequenced a total of 852, 437, 443 and 866 million reads, respectively. The data from CEL-seq consisted of 102 million reads from a total of 74 cells (Figure 1), Supplementary (Figure 1b). All data were processed identically, with cDNA reads clipped to 45 bp, mapped using STAR^27^ and UMIs being quantified using the Drop-seq pipeline^17^. To adjust for differences in sequencing depths, we used only cells with at least one million reads, resulting in 40, 79, 93, 162, 187 cells for CEL-seq, Drop-seq, SCRB-seq, Smart-seq/C1 and Smart-seq2, respectively. To exclude doublets (libraries generated from two or more cells) in the Smart-seq/C1 data, we analyzed microscope images of the microfluidic chips and identified 16 reaction chambers with multiple cells that were excluded from further analysis. For the three UMI methods, we calculated the number of UMIs per library and found that – at least in our case of a rather homogenous cell population – doublets can be readily identified as libraries that have more than twice the mean total UMI count (Supplementary (Figure 1c), which lead to the removal of 0, 3 and 9 cells for CEL-seq, Drop-seq and SCRB-seq, respectively.

Finally, to remove low-quality libraries, we used a method that exploits the fact that transcript detection and abundance in low-quality libraries correlate poorly with high-quality libraries as well as with other low-quality libraries^28^. We therefore determined the maximum Spearman correlation coefficient for each cell in all-to-all comparisons, which readily allowed the identification of low-quality libraries by visual inspection of the distributions of correlation coefficients (Supplementary (Figure 1c). This filtering led to the removal of 5, 16, 30 cells for CEL-seq, Smart-seq/C1, Smart-seq2, respectively, while no cells were removed for Drop-seq and SCRB-seq. The higher number for the two Smart-seq methods is consistent with the notion that in the early barcoding methods (CEL-seq, Drop-seq, SCRB-seq), low-quality cells are probably outcompeted by high-quality cells so that they do not pass our one million reads filter. As Smart-seq/C1 and Smart-seq2 libraries are generated in separate reactions, filtering by correlation coefficient is more important for these methods.

In summary, we processed and filtered our data so that we could use a total of 482 high-quality, equally sequenced scRNA-seq libraries for a fair comparison of the sensitivity, accuracy, precision and efficiency of the methods.

### Single-cell libraries are sequenced to a reasonable level of saturation at one million reads

For all five methods >50% of the reads mapped to the mouse genome (Figure 3a), comparable to previous results^7,16^. Overall, between 48% (Smart-seq2) and 32% (CEL-seq) of all reads were exonic and thus used to quantify gene expression levels. However, the UMI data showed that only 12 %, 5 % and 15 % of the exonic reads were derived from independent mRNA molecules for CEL-seq, Drop-seq and SCRB-seq, respectively (Figure 3a). This indicates that – at the level of mRNA molecules – most of the libraries complexity has already been sequenced at one million reads. To quantify the relationship between the number of detected genes or mRNA molecules and the number of reads in more detail, we downsampled reads to varying depths and estimated to what extend libraries were sequenced to saturation (Supplementary (Figure 2). The number of unique mRNA molecules plateaued at 28,632 UMIs per library for CEL-seq, increased only marginally at 17,207 UMIs per library for Drop-seq and still increased considerably at 49,980 UMIs per library for SCRB-seq (Supplementary (Figure 2c). Notably, CEL-seq showed a steeper slope at low sequencing depths than both Drop-seq and SCRB-seq, potentially due to a less biased amplification by in vitro transcription. Hence, among the UMI methods we found that SCRB-seq libraries had the highest complexity of mRNA molecules that was not yet sequenced to saturation at one million reads. To investigate saturation also for non-UMI-based methods, we applied a similar approach at the gene level by counting the number of genes detected by at least one read. By downsampling, we estimated that ~90% (Drop-seq, SCRB-seq) to 100% (CEL-seq, Smart-seq/C1, Smart-seq2) of all genes present in the library were detected at 1 million reads (Figure 3b), Supplementary (Figure 2a). In particular, the deep sequencing of Smart-seq2 libraries showed clearly that the number of detected genes did not change when increasing the sequencing depth from one million to five million reads per cell (Supplementary (Figure 2b).

**Fig. 3.**
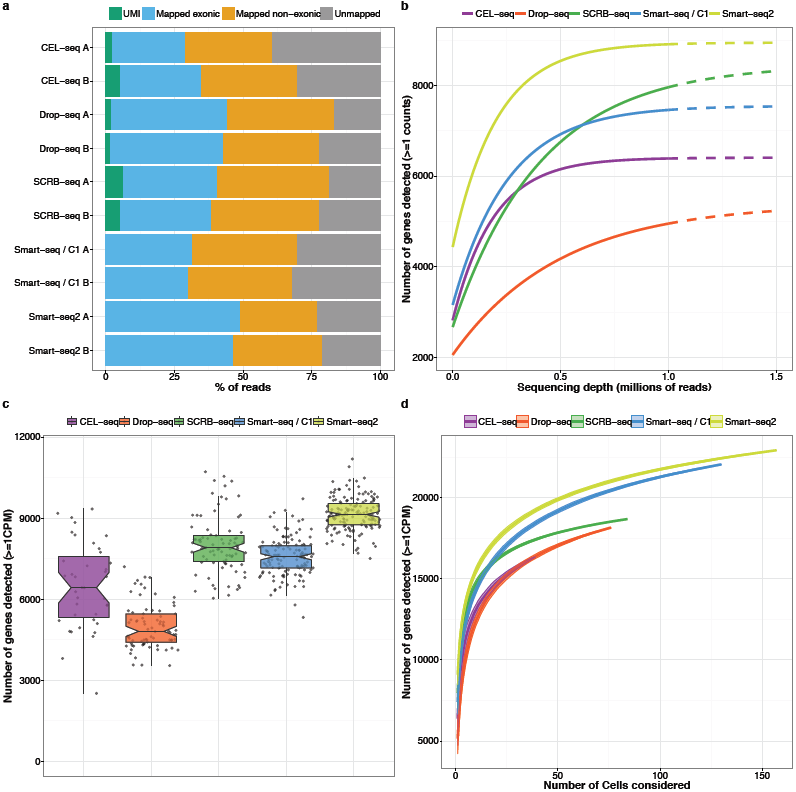
| Sensitivity of scRNA-seq methods. (a) Percentage of reads (downsampled to 1 million per cell) that can not be mapped to the mouse genome (grey), are mapped to regions outside exons (orange), inside exons (blue) and carry a unique UMI (green). For UMI methods, blue denotes the exonic reads amplified from unique UMIs. (b) Median number of genes detected per cell (counts >=1) when downsampling total read counts to the indicated depths. Dashed lines above 1 million reads represent extrapolated asymptotic fits. (c) Number of genes detected (counts >=1) per cell. Each dot represents a cell and each box represents the median, first and third quartile per replicate and method. (d) Cumulative number of genes detected as more cells are added. The order of cells considered was drawn randomly 100 times to display mean ± standard deviation (shaded area).

All in all, these analyses show that single-cell RNA-seq libraries are sequenced to a reasonable level of saturation at one million reads, a cut-off that has also been previously suggested for scRNA-seq datasets^7,29^. While it can be more efficient to analyze scRNA-seq data at lower coverage (see power analyses below), one million reads per cell can be considered as a reasonable starting point for our purpose of comparing scRNA-seq methods.

### Smart-seq2 has the highest sensitivity

Taking the number of detected genes per cell as a measure to compare the sensitivity of the five methods, we found that Drop-seq had the lowest sensitivity with a median of 4811 genes detected per cell, while with CEL-seq, SCRB-seq and Smart-seq/C1 6839, 7906 and 7572 genes per cell were detected, respectively (Figure 3c). Smart-seq2 detected the highest number of genes per cell, with a median of 9138. To compare the total number of genes detected across several cells, we pooled 35 cells per method and detected ~16,000 genes for CEL-seq and Drop-seq, ~17,000 for SCRB-seq, ~18,000 for Smart-seq/C1 and ~19,000 for Smart-seq2 (Figure 3d). While the vast majority of genes (~12,000) were detected by all methods, ~500 genes were specific to each of the 3’ counting methods, but ~1000 genes were specific to each of the two full-length methods (Supplementary (Figure 3a), (Figure 3b). That the full length methods detect more genes in total is also apparent when plotting the genes detected in all available cells, as the 3’ counting methods level off well below 20,000 genes while the two full-length methods level off well above 20,000 genes (Figure 3d).

How evenly reads are distributed across mRNAs can be regarded as another measure of sensitivity. As expected, the 3’ counting methods showed a strong bias of reads mapped to the 3’ end (Supplementary (Figure 4a). However, it is worth mentioning that a considerable fraction of reads also covered more 5’ regions, probably due to internal oligo-dT priming^30^. Smart-seq2 showed a more even coverage than Smart-seq, confirming previous findings^8^. A general difference between the 3’-counting and the full-length methods can also be seen in the quantification of expression levels as they are separated by the first principal component explaining 75% of the total variance (Supplementary (Figure 4b).

**Fig. 4.**
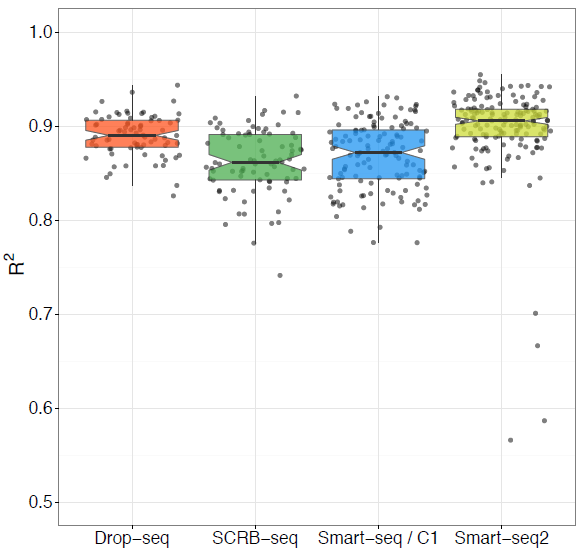
| Accuracy of scRNA-seq methods. ERCC expression values were correlated to their annotated molarity. Shown are the distributions of correlation coefficients (adjusted R^2^ of linear regression model) across methods. Each dot represents a cell/bead and each box represents the median, first and third quartile.

As an absolute measure of sensitivity, we compared the probability of detecting the 92 spiked-in ERCCs, for which the number of molecules available for library construction is known (Supplementary (Figure 5). We determined the detection probability of each ERCC mRNA as the proportion of cells with non-zero read or UMI counts^31^. For the CEL-seq data, Gruen et al. noted that their ERCCs were likely degraded^23^ and we also found that ERCCs from the CEL-seq data are detected with a ten-fold lower efficiency than for the other methods (data not shown). Therefore, we did not consider the CEL-seq libraries for any ERCC-based analyses. For Drop-seq, we used the ERCC-only data set^17^ and for the other three methods, 2-5% of the one million reads per cell mapped to ERCCs, which were sequenced to complete saturation at that level (Supplementary (Figure 5b). For Smart-seq2, an ERCC RNA molecule was detected on average in half of the libraries when ~7 molecules were present in the sample, while Smart-seq/C1 required ~11 molecules for detection in half of the libraries. Drop-seq and SCRB-seq has estimates of ~16-17 molecules per cell (Supplementary (Figure 5c)–(e).

**Fig. 5.**
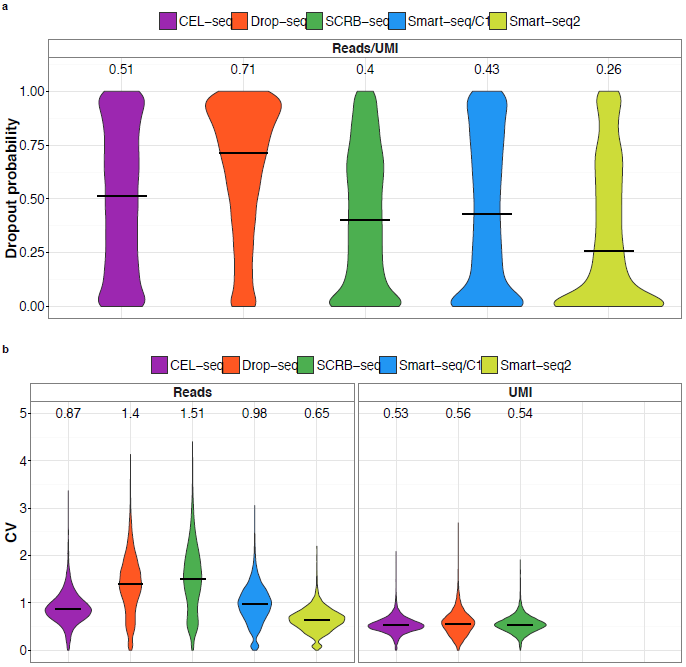
| Precision of scRNA-seq methods. We compared precision among methods using the 12,942 genes detected in at least 25% of all cells by any method in a subsample of 35 cells per method. (a) Distributions of dropout rates across the 12,942 genes are shown as violin plots and medians are shown as bars and numbers. (b) Distributions of the coefficient of variation (CV) across the 12,942 genes calculated from cells with at least one count are shown as violin plots and medians are shown as bars and numbers. For 1096, 1487, 480, 904, 621 genes for CEL-seq, Drop-seq, SCRB-seq, Smart-seq/C1 and Smart-seq2, respectively no CV could be calculated as fewer than two cells had non-zero counts. Including these genes with a high CV would result in median values of 0.9/0.54, 1.49/0.6, 1.54/0.55, 1.01, 0.66, respectively.

Notably, the sensitivity estimated from the number of detected genes does not fully agree with the comparison based on ERCCs. While Smart-seq2 is the most sensitive method in both cases, Drop-seq performs better and SCRB-seq performs worse when using ERCCs. The reasons for this discrepancy are unclear, but several have been noted before^32–34^ including that ERCCs do not model endogenous mRNAs perfectly since they are shorter, have shorter poly-A tails, lack a 5’ cap and can show batch-wise variation in concentrations as observed for the CEL-seq data. In the case of Drop-seq, it should be kept in mind that ERCCs were sequenced separately as discussed above and in this way leading to a higher efficiency. Therefore, while it is still useful to estimate the absolute range in which molecules are detected, for our purpose of comparing the sensitivity of methods using the same cells, we regard the number of detected genes per cell as the more reliable estimate of sensitivity in our setting, as it sums over many, non-artificial genes.

In summary, we find that Smart-seq2 is the most sensitive method as it detects the highest number genes per cell, the most genes in total across cells and has the most even coverage of transcripts. Smart-seq/C1 is slightly less sensitive per cell, but detects the same number of genes across cells, if one considers its lower fraction of mapped exonic reads (Figure 3a). Among the 3’ counting methods, SCRB-seq is most sensitive, closely followed by CEL-seq, whereas Drop-seq detects considerably fewer genes.

### Accuracy is similar across scRNA-seq methods

In order to quantify the accuracy of transcript level quantifications, we compared observed expression values with annotated molecule concentration of the 92 ERCC transcripts (Supplementary (Figure 5a). For each cell, we calculated the correlation coefficient (R^2^) for a linear model fit (Figure 4). The median accuracy did differ among methods (Kruskal-Wallis test, p<2.2e-16) with Smart-seq2 having the highest accuracy, especially since it is more accurate at lower concentrations (Supplementary (Figure 6). Importantly, all methods had fairly high accuracies ranging between 0.86 and 0.91, suggesting that they all measure absolute mRNA levels fairly well. As discussed above, CEL-seq was excluded from the ERCC analyses due to the potential degradation of the ERCCs in this data set^23^. The original publication for CEL-seq from 10 pg of total RNA input and ERCC spike-in reported a mean correlation coefficient of R^2^=0.87^18^, similar to the correlations reported for the other four methods. Hence, we find that the accuracy is similarly high across all five methods and also because absolute expression levels are rarely of interest, the small differences in accuracy will rarely be a decisive factor when choosing among the five methods.

**Fig. 6.**
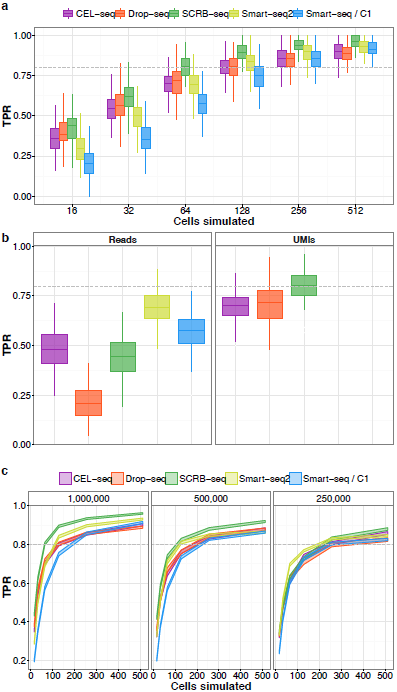
| Power of scRNA-seq methods. Using the empirical mean/dispersion and mean/dropout relationships (Supplementary Fig. 10), we simulated data for two groups of *n* cells each for which 5% of the 12,942 genes were differentially expressed with log-fold changes drawn from observed differences between microglial subpopulations from a previously published dataset^37^. The simulated data were then tested for differential expression using limma^38^, from which the average true positive rate (TPR) and the average false discovery rate (FDR) was calculated (Supplementary Fig. 12). (a) TPR for 1 million reads per cell for sample sizes n=16, n=32, n=64, n=128, n=256 and n=512 per group. Boxplots represent the median, first and third quartile of 100 simulations. (b) TPR for 1 million reads per cell for n=64 per group with and without using UMI information. Boxplots represent the median, first and third quartile of 100 simulations. (c) TPRs as in (a) using mean/dispersion and mean/dropout estimates from 1 million (as in (a)), 0.5 million and 0.25 million reads. Line areas indicate the median power with standard error from 100 simulations..

### Precision is determined by a combination of dropout rates and amplification noise and is highest for SCRB-seq

While a high accuracy is necessary to quantify absolute expression values, one of the most common experimental aims is to compare relative expression levels in order to identify differentially expressed genes or biological variation among cells. Hence, the precision, i.e. the reproducibility or the amount of technical variation -is the major factor of a method. As we used the same cells under the same culture conditions, we assume that the amount of biological variation is the same across all five methods. Hence, all differences in the total variation between methods are due to technical variation. Technical variation is substantial in scRNA-seq data because a substantial fraction of mRNAs is lost during cDNA generation and small amounts of cDNA get amplified. Therefore, both these processes, the dropout probability and the amplification noise, need to be considered when quantifying variation. Indeed, a mixture model including a dropout-rate, and a negative binomial distribution, modelling the overdispersion in the count data, have been shown to represent scRNA-seq data better than the negative binomial alone^35,36^.

To compare the methods for a common set of genes without penalizing more sensitive methods, we selected the 12,942 genes that were detected in 25% of the cells by at least one method (Supplementary Fig. 7). We then assessed their technical variation in a subsample of 35 cells per method to exclude any bias due to the different numbers of cells analysed in each method. We measured the loss of molecules in cDNA generation as the fraction of cells with zero counts (Fig. 5a, Supplementary Fig. 7b). As expected from the number of detected genes per cell (Fig. 3c), Drop-seq had the highest median dropout probability (71%) and Smart-seq2 the lowest (26%). To assess the variation due to the amplification of the detected genes, we calculated the coefficient of variation (CV, standard deviation divided by the mean) of all cells with non-zero counts. As expected from the removal of amplification noise when using UMI information, the three UMI methods showed the lowest median variation, even when considering that the CV could not be calculated for all genes (Figure 5b). When ignoring UMI information, it becomes apparent that Smart-seq2 is the protocol that has the lowest amplification noise and that the reduction in amplification noise due to UMIs is considerable (Figure 5b), Supplementary Fig. 9). The latter effect has been previously described for CEL-seq^23^ and is even stronger for SCRB-seq and Drop-seq, fitting with the notion that in vitro amplification is more precise than PCR amplification. In summary, Smart-seq2 measures the common set of 12,942 in more cells with more amplification noise and the UMI methods measure them in fewer cells with less amplification noise. However, how the different combinations of dropout rates and amplification precisions affect the power to detect e.g. differentially expressed genes is not evident, neither from this analysis nor from variation measures that combine dropout probabilities and amplification precision (CV of all cells, Supplementary Fig. 8 and 9).

Therefore, we conducted power simulations that used for each method the observed mean-variance and mean-dropout relationship for the 12,942 genes. First, we estimated the mean and dispersion parameter (i.e. the shape parameter of the gamma mixing distribution) for each gene per method. Next, we fitted a spline to the resulting pairs of mean and dispersion estimates in order to predict the dispersion of a gene given its mean (Supplementary Fig. 10a). Finally, we included the sensitivity of each scRNA-seq method in the power simulations by modeling a gene-wise dropout parameter from the observed detection rates also dependent on the mean expression (Supplementary Fig. 10b). When simulating data according to these fits, we recovered distributions of dropout rates and amplification noise closely matching the observed data (Supplementary Fig. 11). To compare the power for differential gene expression among the methods, we simulated read counts for two groups of cells by adding log-fold changes to 5% of the genes. These logfold changes were drawn from observed differences between microglial subpopulations from a previously published dataset^37^ to mimic a biologically realistic scenario. The simulated datasets were then tested for differential expression using limma^38^, from which the average true positive rate (TPR) and the average false discovery rate (FDR) could be calculated for all the 12,942 genes.

First, we analyzed how the number of analyzed cells affects TPR and FDR by running 100 simulations each for a range of 16 to 512 cells per group. SCRB-seq performed best, reaching a median TPR of 80% with 64 cells. CEL-seq, Drop-seq and Smart-seq2 performed slightly worse, reaching 80% power with 103, 99 and 95 cells per group, respectively, while Smart-seq/C1 needed with 150 cells per group considerably more cells to reach 80% power (Figure 6a). FDRs were similar in all methods and just slightly above 5% (Supplementary Fig. 12). As expected from the effect of UMIs on amplification noise (Figure 5b), Smart-seq2 performed best when just considering reads and UMIs strongly increased the power, especially for Drop-seq and SCRB-seq (Figure 6b).

Next, we asked how TPR and FDR depend on the sequencing depth. We repeated our simulation studies as described above, but estimated the mean-dispersion and mean-dropout relationships from data downsampled to 500,000 or 250,000 reads per cell. Overall, the decrease in power was moderate (Figure 6c), Table 1) and mainly linked to decreased gene detection rates. Importantly, not all methods responded to downsampling at similar rates, congruent with their different relationship of sequenced reads and detected genes (Supplementary (Figure 2). While Smart-seq2 was only slightly affected and reached 80% power with 95, 105 and 128 cells at 1, 0.5 and 0.25 million reads, respectively, SCRB-seq and Drop-seq required 2.6 fold more cells at 0.25 million reads than at 1 million reads (Table 1). In summary, at one million reads and half a million reads SCRB-seq is the most precise, i.e. most powerful method, but at a sequencing depth of 250,000 reads Smart-seq2 needs the lowest number of cells to reach 80% power. The optimal balance between the number of cells and their sequencing depth depends on many factors, but the monetary cost is certainly an important one. Hence, we used the results of our simulations to compare the costs among the methods for a given level of power.

### Cost-Efficiency is similarly high for drop-seq, SCRB-seq and smart-seq2

Given the number of single cells that are needed per group to reach 80% power as simulated above for three sequencing depths (Figure 6c), we calculated the minimal costs to generate and sequence these libraries. For example, at one million reads, SCRB-seq requires 64 cells per group. Generating 128 SCRB-seq libraries costs ~260€ and generating 128 million reads costs ~640€. Note, that the necessary paired-end reads for CEL-seq, SCRB-seq and Drop-seq can be done with a 50 cycles single end kit and hence we assume that sequencing costs are the same for all methods.

**Table 1.** Cost efficiency extrapolation for single-cell RNA.seq experiments.

| Method | TPR<sup>a</sup> | FDR<sup>a</sup> (%) | Cell per group<sup>b</sup> | Library cost (€) | Minimal cost<sup>c</sup> (€) |
| --- | --- | --- | --- | --- | --- |
| CEL-seq | 0.8 | ~5.7 | 103 \| 138 \| 181 | ~8 | ~ 2680 \| 2900 \| 3350 |
| Drop-seq | 0.8 | ~8.4 | 99 \| 135 \| 254 | ~0.1 | ~ 1010 \| 700 \| 690 |
| SCRB-seq | 0.8 | ~6.1 | 64 \| 90 \| 166 | ~2 | ~ 900 \| 810 \| 1080 |
| Smart-seq/C1 | 0.8 | ~4.9 | 150 \| 172 \| 215 | ~25 | ~ 9010 \| 9440 \| 11290 |
| Smart-seq2<br>(commercial) | 0.8 | ~5.1 | 95 \| 105 \| 128 | ~30 | ~ 10470 \| 11040 \| 13160 |
| Smart-seq2<br>(in-house Tn5) | 0.8 | ~5.1 | 95 \| 105 \| 128 | ~3 | ~ 1520 \| 1160 \| 1090 |

<sup>a</sup> True positive rate and false discovery rate based on simulations (Fig. 6 and Supplementary Fig. 12);
<sup>b</sup> sequencing depth of 1, 0.5 and 0.25 million reads; <sup>c</sup> Assuming 5 € per one million reads;

When we do analogous calculations for the four other methods, Drop-seq is with 690 € the most cost-efficient method when sequencing 254 cells at a depth of 250,000 reads (Table 1, Supplementary Fig. 13). SCRB-seq is just slightly more expensive in this calculation and also Smart-seq2 is with 1090 € in the same range, as long as one uses in-house Tn5 transposase^26^ as was also done in our experiments. When using the commercial Nextera kit as described^9^, the costs for Smart-seq2 are ten-fold higher and even if one reduces the amount of Nextera transposase as in the Smart-seq/C1 protocol by 4-fold, the published Smart-seq2 protocol is four times more expensive than the early barcoding methods. Smart-seq/C1 is almost ten-fold less efficient due its high library costs that arise from the microfluidic chips, the commercial Smart-seq kit and the costs for commercial Nextera XT kits.

Of note, these calculations are the minimal costs of the experiment and several factors are not considered such as costs to set-up the methods, costs to isolate single cells or costs due to practical constraints in generating a fixed number of scRNA-seq libraries. In particular, it is important that independent biological replicates are needed when investigating particular factors such as genotypes or developmental timepoints and Smart-seq/C1 and Drop-seq are less flexible in distributing scRNA-seq libraries across replicates than the other three methods that use PCR-plates. Furthermore, the costs are increased by unequal sampling from the included cells as well as from sequencing reads from cells that are excluded. In our case, between 6% (CEL-seq, SCRB-seq) and 32% (Drop-seq) of the reads came from cell barcodes that were not included. While it is difficult to accurately and transparently compare these costs among the methods, it is evident that they will increase the costs for Drop-seq relatively more than for the other methods. In summary, we find that Drop-seq, SCRB-seq are the most cost-efficient methods, closely followed by Smart-seq2 when producing one's own transposase.

## Discussion

Single-cell RNA-sequencing (scRNA-seq) is a powerful technology to tackle a multitude of biomedical questions. To facilitate choosing among the many approaches that were recently developed, we systematically compared five scRNA-seq methods and assessed their sensitivity, accuracy, precision and cost-efficiency. We chose a leading commercial platform (Smart-seq/C1), one of the most popular full-length methods (Smart-seq2), a UMI-method that uses in-vitro transcription for cDNA amplification from manually isolated cells (CEL-seq), a UMI-method that has a very high throughput (Drop-seq) and a UMI-method that allows single-cell isolation by FACS (SCRB-seq). Protocols are available for all these methods and can therefore be set up by any molecular biology lab.

We find that SCRB-seq, Smart-seq/C1 and CEL-seq detect a similar number of genes per cell, while Drop-seq detects nearly 50% less than the most sensitive method Smart-seq2 (Figure 3b),(Figure 3c). Despite this lower per cell sensitivity, Drop-seq does not generally detect fewer genes since the total number of detected genes converges around 18,000, similar as for SCRB-seq and CEL-seq (Fig. 3d). A potential explanation could be that a fraction of mRNA molecules gets randomly detached from the beads when droplets are broken up for reverse transcription. It will be interesting to see whether this step could be optimized in the future. While the three 3’ counting methods detect largely the same set of genes, Smart-seq/C1 and Smart-seq2 detect around 3000 additional genes (Figure 3d), Supplementary Fig. 3b), suggesting that some 3’ ends of cDNAs might be difficult to convert to sequenceable molecules. When using ERCCs to compare absolute sensitivities, we again find Smart-seq2 to be the most sensitive method. However, we also find that sensitivity estimates from ERCCs do not perfectly correlate with estimates from endogenous genes, suggesting that they might not always be an ideal benchmark for comparing different methods. In summary, we find that Smart-seq2 is the most sensitive method based on its gene detection rate per cell and in total. In addition, Smart-seq2 shows the most even read coverage across transcripts (Supplementary Fig. 4a), making it the most appropriate method for detecting alternative splice forms. Hence, Smart-seq2 is certainly the most suitable method when an annotation of single cell transcriptomes is the focus.

We find that Smart-seq2 is the also the most accurate method, i.e. has the highest correlation of known mRNA concentrations and read counts per million. Importantly, accuracy is similarly high across all methods. As absolute quantification of mRNA molecules is rarely of interest, the slight differences in accuracy will rarely be an important criterion for choosing among the five methods. In contrast, relative quantification of gene expression levels is of interest for most scRNA-seq studies and hence the precision of the method is an important benchmark.

How precisely a gene is measured depends on two factors: in how many cells it is measured and, if it is measured, how reproducibly this is done. For the first factor (dropout probability), we find Smart-seq2 to be the best method (Figure 5a), as expected from its highest gene detection sensitivity. For the second factor (CV of non-zero cells), we find the three UMI methods to perform best (Figure 5b), as expected from their ability to eliminate variation introduced by amplification^21^. To assess the importance of these two factors in combination, we performed simulations and could show that SCRB-seq has the highest power to detect differentially expressed genes (Figure 6). This strongly depends on the use of UMIs, especially for the PCR-based amplification methods (Figure 5b) and (Figure 6b) and the strong reduction in amplification noise can also be seen in mean-variance plots (Supplementary Fig. 9). Notably, this is due to the large amount of amplification needed for scRNA-seq libraries, as the effect of UMIs on power for bulk RNA-seq libraries is neglectable^39^.

Maybe practically most important, our power simulations also allow to compare the efficiency of the methods by calculating the costs to generate the data for a given level of power. Using simple, minimal cost calculations we find that Drop-seq is the most efficient method, closely followed by SCRB-seq and Smart-seq2. However, when considering that Drop-seq costs are more underestimated than those for SCRB-seq and Smart-seq2, due to its lower flexibility in generating a fixed number of libraries and due to its higher fraction of reads that come from cells that are excluded, Drop-seq and SCRB-seq are in practice probably similar efficient. For Smart-seq2 to be similar efficient it is absolutely necessary to use in-house produced transposase as described^26^ and also done here.

Given comparable efficiencies of Drop-seq, SCRB-seq and Smart-seq2, additional factors will play a role when choosing a suitable method for a particular question. Due to its low library costs, Drop-seq is probably preferable when analysing large numbers of cells at low coverage, e.g. to find rare cell types. On the other hand, Drop-seq in its current setup requires a relatively large amount of cells (>6,500 for one minute of flow). Hence, if few and/or unstable cells are isolated by FACS, SCRB-seq or Smart-seq2 is probably preferable. Additional advantages of these two methods over Drop-seq include that technical variation can be estimated from ERCCs for each cell, which is helpful to estimate biological variation^40–42^, and that the exact same setup can be used to generate bulk RNA-seq libraries. While SCRB-seq is slightly more efficient than Smart-seq2 and has the advantage that one does not need to produce transposase in-house, Smart-seq2 is preferable when transcriptome annotation and the quantification of different splice forms are of interest. So while such factors will be differently weighted by each individual lab and for each research question, our analyses provide a solid basis for such considerations when choosing among the analysed methods.

While we find comparable efficiencies for Drop-seq, SCRB-seq and Smart-seq2, CEL-seq and even more so Smart-seq/C1 and Smart-seq2 (using commercial transposase) are 313-fold less efficient and cannot compete at these precisions and costs. However, a CEL-seq2 protocol with considerable improvements in sensitivity and costs has recently been published^43^. The sensitivity of this CEL-seq2 protocol is even further improved when run in the nanoliter volumes of the C1 device^43^, again showing that small volumes in general improves sensitivity^7^. Also the efficiency of the Fluidigm C1 platform can be further increased by microfluidic chips with a higher throughput, as available in the HT mRNA-seq IFC. Our finding that Smart-seq2 is the most sensitive protocol when ignoring UMIs, also hints towards further possible improvements of SCRB-seq and Drop-seq. As these methods also rely on template switching and PCR amplification, the improvements found in the systematic optimization of Smart-seq2^8^ could also improve the sensitivity of SCRB-seq and Drop-seq. Furthermore, the costs of SCRB-seq libraries per cell can be halfed when switching to a 384-well format^15^. Hence, it is clear that scRNA-seq protocols can and will be further improved and that our analysis does not provide a final answer on which scRNA-seq method is most efficient. However, our analysis does allow an informed choice for five prominent current scRNA-seq methods and – maybe more importantly – provides a framework and starting point for comparative evaluations in the future.

## Conclusions

We systematically compared five prominent scRNA-seq methods and find that Drop-seq is preferable when quantifying transcriptomes of a large numbers of cells with low sequencing depth, SCRB-seq is preferable when quantifying transcriptomes of fewer cells and Smart-seq2 is preferable when annotating and/or quantifying transcriptomes of fewer cells as long one can use in-house produced transposase. Our analysis allows an informed choice among the tested methods and provides a solid framework for benchmarking future improvements in scRNA-seq methodologies.

## Methods

### Published data

CEL-seq data for J1 mESC cultured in 2i/LIF condition^23^ were obtained under accession GSE54695. Drop-seq ERCC^17^ data were obtained under accession GSE66694. Raw fastq files were extracted using the SRA toolkit (2.3.5). We trimmed cDNA reads to the same length and processed raw reads in the same way as data sequenced for this study.

### Cell culture of mESC

J1 mouse embryonic stem cells were maintained on gelatin-coated dishes in Dulbecco's modified Eagle's medium supplemented with 16% fetal bovine serum (FBS, Sigma-Aldrich), 0. 1 mM β-mercaptoethanol (Invitrogen), 2 mM L-glutamine, 1x MEM non-essential amino acids, 100 U/ml penicillin, 100 μg/ml streptomycin (PAA Laboratories GmbH), 1000 μ/ml recombinant mouse LIF (Millipore) and 2i (1 μM PD032591 and 3 μM CHIR99021 (Axon Medchem, Netherlands). J1 embryonic stem cells were obtained from E. Li and T. Chen and mycoplasma free determined by a PCR-based test. Cell line authentication was not recently performed.

### Single cell RNA-seq library preparations

#### Drop-seq

Drop-seq experiments were performed as published^17^ and successful establishment of the method in our lab was confirmed by a species-mixing experiment (Supplementary Fig. 1a). For this work, J1 mES cells (100/μl) and barcode-beads (120/μl, Chemgenes) were co-flown in Drop-seq PDMS devices (Nanoshift) at rates of 4000 |μl/hr. Collected emulsions were broken by addition of perfluoroctanol (Sigma-Aldrich) and mRNA on beads was reverse transcribed (Maxima RT, Thermo Fisher). Unused primers were degraded by addition of Exonuclease I (New England Biolabs). Washed beads were counted and aliquoted for preamplification (2000 beads / reaction). Nextera XT libraries were constructed from 1 ng of pre-amplified cDNA with a custom P5 primer (IDT).

#### SCRB-seq

RNA was stabilized by resuspending cells in RNAprotect Cell Reagent (Qiagen) and RNAse inhibitors (Promega). Prior to FACS sorting, cells were diluted in PBS (Invitrogen). Single cells were sorted into 5 μl lysis buffer consisting of a 1/500 dilution of Phusion HF buffer (New England Biolabs) and ERCC spike-ins (Ambion), spun down and frozen at −80°C. Plates were thawed and libraries prepared as described previously^15^. Briefly, RNA was desiccated after protein digestion by Proteinase K (Ambion). RNA was reverse transcribed using barcoded oligo-dT primers (IDT) and products pooled and concentrated. Unincorporated barcode primers were digested using Exonuclease I (New England Biolabs). Pre-amplification of cDNA pools were done with the KAPA HiFi HotStart polymerase (KAPA Biosystems). Nextera XT libraries were constructed from 1 ng of preamplified cDNA with a custom P5 primer (IDT).

#### Smart-seq/CI

Smart-seq/C1 libraries were prepared on the Fluidigm C1 system according to the manufacturer's protocol. Cells were loaded on a 10-17 μm RNA-seq microfluidic IFC at a concentration of 200,000/ml. Capture site occupancy was surveyed using the Operetta (Perkin Elmer) automated imaging platform.

#### Smart-seq2

mESCs were sorted into 96-well PCR plates containing 2 μl lysis buffer (1.9 μl 0.2% TritonX-100; 0.1 μl RNAseq inhibitor (Lucigen)) and spike-in RNAs (Ambion), spun down and frozen at −80 °C. To generate Smart-seq2 libraries, priming buffer mix containing dNTPs and oligo-dT primers was added to the cell lysate and denatured at 72 °C. cDNA synthesis and pre-amplification of cDNA was performed as described previously^8,9^. Sequencing libraries were constructed from 2.5 ng of pre-amplified cDNA using an inhouse generated Tn5 transposase^26^. Briefly, 5 μl cDNA was incubated with 15 μl tagmentation mix (1 μl of Tn5; 2 μl 10x TAPS MgCl_2_ Tagmentation buffer; 5 μl 40% PEG8000; 7 μl water) for 8 min at 55 °C. Tn5 was inactivated and released from the DNA by the addition of 5 μl 0.2% SDS and 5 min incubation at room temperature. Sequencing library amplification was performed using 5 μl Nextera XT Index primers (Illumina) that had been first diluted 1:5 in water and 15 μl PCR mix (1 μl KAPA HiFi DNA polymerase (KAPA Biosystems); 10μl 5x KAPA HiFi buffer; 1.5 μl 10mM dNTPs; 2.5μl water) in 10 PCR cycles. Barcoded libraries were purified and pooled at equimolar ratios.

### DNA sequencing

For SCRB-seq and Drop-seq, final library pools were size-selected on 2% E-Gel Agarose EX Gels (Invitrogen) by excising a range of 300-800 bp and extracting DNA using the MinElute Kit (Qiagen) according to the manufacturer's protocol.

Smart-seq/C1, Drop-seq and SCRB-seq library pools were sequenced on an Illumina HiSeq1500 using the High Output mode. Smart-seq2 pools were sequenced on Illumina HiSeq2500 (Replicate A) and HiSeq2000 (Replicate B) platforms. Smart-seq/C1 and Smart-seq2 libraries were sequenced 45 cycles single-end, whereas Drop-seq and SCRB-seq libraries were sequenced paired-end with 20 cycles to decode cell barcodes and UMI from read 1 and 45 cycles into the cDNA fragment. Similar sequencing qualities were confirmed by FastQC v0.10.1 (Supplementary Fig. 1b).

### Basic data processing and sequence alignment

Smart-seq/C1/Smart-seq2 libraries (i5 and i7) and Drop-seq/SCRB-seq pools (i7) were demultiplexed from the Nextera barcodes using deML^44^. All reads were trimmed to the same length of 45 bp by cutadapt^45^ and mapped to the mouse genome (mm10) including mitochondrial genome sequences and unassigned scaffolds concatenated with the ERCC spike-in reference. Alignments were calculated using STAR 2.4.0^27^ using all default parameters.

For libraries containing UMIs, cell-and gene-wise count/UMI tables were generated using the published Drop-seq pipeline (v1.0)^17^. We discarded the last 2 bases of the Drop-seq cell and molecular barcodes to account for bead synthesis errors.

For Smart-seq/C1 and Smart-seq2, features were assigned and counted using the Rsubread package (v1.20.2)^46^.

### Power simulations

We developed a framework in R for statistical power evaluation of differential gene expression in single cells. For each method, we estimated the mean expression, dispersion and dropout probability per gene from the same number of cells per method. In the read count simulations, we followed the framework proposed in Polyester^47^, i.e. we retained the observed mean-variance dependency by applying a cubic smoothing spline fit. Furthermore, we included a local polynomial regression fit for the mean-dropout relationship to capture the heteroscedasticity observed. In each iteration, we simulated count measurements for the 12,942 genes for sample sizes of 2^4^, 2^5^, 2^6^, 2^7^, 2^8^ and 2^9^ cells per group. The read count for a gene *i* in a cell *j* is modeled as a product of a binomial and negative binomial distribution:

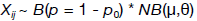

The mean expression magnitude μ was randomly drawn from the empirical distribution. 5 percent of the genes were defined as differentially expressed with an effect size drawn from the observed fold changes between microglial subpopulations in Zeisel et al^37^. The dispersion θ and dropout probability p_θ_ were predicted by above mentioned fits.

For each method and sample size, 100 RNA-seq experiments were simulated and tested for differential expression using limma^38^ in combination with voom^48^ (v3.26.7).

The power simulation framework was implemented in R and is available in Additional File 1, including an example dataset.

### ERCC capture efficiency

To estimate the single molecule capture efficiency, we assume that the success or failure of detecting an ERCC is a binomial process, as described before^31^. Detections are independent from each other and are thus regarded as independent Bernoulli trials. We recorded the number of cells with nonzero and zero read or UMI counts for each ERCC per method and applied a maximum likelihood estimation to fit the probability of successful detection. The fit line was shaded with the 95% Wilson score confidence interval.

### Cost efficiency calculation

We based our cost efficiency extrapolation on the power simulations starting from empirical data at different sequencing depths (250,000 reads, 500,000 reads, 1,000,000 reads; (Figure 6c). We determined the number of cells required per method and depth for adequate power (80%) by an asymptotic fit to the median powers. For the calculation of sequencing cost, we assumed 5€ per million raw reads, independent of method. Although UMI-based methods need paired-end sequencing, we assumed a 50 cycle sequencing kit is sufficient for all methods.

## Declarations

### Ethics approval and consent to participate

Not applicable

### Consent for publication

Not applicable

### Availability of data and material

The raw and analyzed data files can be obtained in GEO under accession number GSE75790.

## Author's contributions

CZ and WE conceived the experiments. CZ prepared scRNA-seq libraries and analyzed the data. BV implemented the power simulation framework and estimated ERCC capture efficiencies. SP helped in data processing and power simulations. BR prepared Smart-seq2 scRNA-seq libraries. MS performed cell culture of mESC. WE and HL supervised the experimental work and IH provided guidance in data analysis. CZ, IH, BR and WE wrote the manuscript. All authors read and approved the final manuscript.

## Acknowledgements

We thank Rickard Sandberg for facilitating the Smart-seq2 sequencing and fruitful discussion. We thank Christopher Mulholland for assistance with FACS sorting, Dominik Alterauge for help establishing the Drop-seq method and Stefan Krebs and Helmut Blum from the LAFUGA platform for sequencing. We are grateful to Magali Soumillon and Tarjei Mikkelsen for providing the SCRB-seq protocol.

